# High-resolution spatial analysis reveals pregnenolone metabolism as a potential target for immuno-suppressive microenvironment of *BRAF* mutant colorectal cancer

**DOI:** 10.1101/2025.09.01.673435

**Authors:** Xicheng Wang, Zihao Zhao, Lian Liu, Jie Cheng, Yi Ba

**Affiliations:** Cancer Medical Center, Peking Union Medical College Hospital, Chinese Academy of Medical Sciences, Beijing, China; Department of pathology, School of Basic Medicine, Tongji Medical College and State Key Laboratory for Diagnosis and Treatment of Severe Zoonotic Infectious Diseases, Huazhong University of Science and Technology, Hubei, China; Institute of Pathology, Tongji Hospital, Huazhong University of Science and Technology, Hubei, China; School of Life Sciences, Center for Statistical Science, Peking University, Beijing, China

**Author notes:** Correspondence: Y.B., J.C. These authors contribute equally (Xicheng Wang, Zihao Zhao).

**Keywords:** BRAF mutant colorectal cancer, pregnenolone, CYP11A1, CD8^+^ T cell exhaustion, spatial transcriptomics, immunotherapy

## Abstract

*BRAF* mutant colorectal cancer (CRC) is highly aggressive and often resistant to traditional therapies, presenting a significant challenge in cancer immunotherapy. Understanding the tumor microenvironment (TME) and identifying novel immune checkpoints are crucial for improving treatment outcomes. Here, we employed high-resolution spatial analysis and single-cell RNA sequencing to map the TME of *BRAF* mutant CRC at single-cell resolution, focusing on the interactions between exhausted CD8^+^ T cells and C1QC^+^ macrophages. Mechanistically, we discovered that C1QC^+^ macrophages drive CD8^+^ T cell exhaustion through pregnenolone synthesis via the steroidogenic enzyme CYP11A1. Our findings identify pregnenolone metabolism as a previously unrecognized immunosuppressive pathway in *BRAF* mutant CRC, and suggest that steroid biosynthesis is a generalizable mechanism of immune evasion beyond endocrine-related malignancies. Targeting this pathway offers a promising therapeutic strategy for enhancing anti-tumor immunity in *BRAF* mutant CRC and potentially other cancers.

## Introduction

Colorectal cancer (CRC) ranks second worldwide in terms of mortality in both sexes [1]. *BRAF* gene mutations occur in 7% of patients with CRC [2], of which approximately 90% have *BRAF* V600E mutations [3]. *BRAF* mutant advanced CRC is highly aggressive and is associated with worse clinical outcomes [4]. *BRAF* mutant CRC exhibit an approximately 70% increase in mortality compared to *BRAF* wild-type CRC [5]. The MEK116833 study (NCT01750918) and the BEACON study (NCT02928224) showed that the objective response rates (ORRs) of anti-EGFR monoclonal antibodies in combination with *BRAF* inhibitors as second-line treatment of *BRAF* mutant advanced CRC were only 10% and 20%, respectively, and the median progression-free survival (PFS) from the two trials was 3.5 months and 4 months [6, 7], respectively. This phenomenon illustrates that disease progression and drug resistance are inevitable in *BRAF* mutant CRC. However, research on the drug resistance mechanism of *BRAF* mutant advanced CRC remains limited [8, 9]. Therefore, it is important to further investigate the mechanism of drug resistance in *BRAF* mutant CRC patients and to find new therapeutic targets in the TME to overcome this resistance.

The immune microenvironment of *BRAF* mutant cancer is characterized by distinct immunosuppressive features that contribute to poor prognosis and resistance to therapies including immune checkpoint inhibitors (ICIs). *BRAF* mutations lead to dysregulated signaling in the MAPK pathway and shape the tumor immune landscape [10, 11]. However, the key features of the immune microenvironment in *BRAF* mutant CRC remain unknown. Both single-cell and spatial transcriptomic landscapes remain unexplored, which hinders mechanistic research on immunotherapy for *BRAF*-mutant CRC. Understanding the immune microenvironment of *BRAF* mutant CRC provides critical insights into therapeutic resistance mechanisms and highlights novel strategies to improve treatment outcomes. Herein, thoroughly exploring the immune cell composition and intercellular interactions within the immune microenvironment may help demonstrate the immune profiles and cellular processes of *BRAF* V600E mutant CRC patients.

Steroid hormones, such as pregnenolone and its derivatives, have long been studied for their roles in hormone-dependent cancers, such as prostate and breast cancers. However, their functions in non-hormone-related tumors remain poorly understood. Pregnenolone, which is synthesized from cholesterol by the mitochondrial enzyme CYP11A1, is a precursor of all steroid hormones and has been shown to modulate immune responses and cancer progression in prostate cancer and melanoma. Despite this, little attention has been paid to its role in colorectal cancer.

Here, we elucidated the TME in *BRAF* mutant CRC patients using single-cell mapping and spatial mapping with single-cell precision. We found that the interaction between exhausted CD8^+^ T cells and C1QC^+^ macrophages plays a key role in the immunosuppressive tumor microenvironment of *BRAF* V600E mutant CRC. In particular, C1QC^+^ macrophage-derived pregnenolone promoted immune evasion by inducing the exhaustion of CD8^+^ T cells. Our findings suggest that pregnenolone, produced by tumor-associated macrophages, contributes to CD8^+^ T cell exhaustion and immune evasion. This extends the oncological relevance of steroidogenesis beyond classic hormone-responsive tumors and introduces pregnenolone metabolism as a novel and targetable axis in immunotherapy for colorectal cancer.

## Results

### Microenvironment of *BRAF* mutant CRC is highly immune suppressive

To elucidate the microenvironment of *BRAF* mutant CRC, we collected published scRNA-seq data of CRC and classified them into wild-type and *BRAF* mutant groups according to WES data. After filtering the scRNA-seq data to exclude damaged or dead cells and putative doublets, 191,292 single-cell transcriptomes from 50 patients were retained for subsequent analyses. A total of 176,856 cells were obtained from *BRAF* wild-type (WT) tumors, and 14,436 cells were obtained from *BRAF*-mutant (MUT) tumors. Following the normalization of gene expression based on sequencing depth and mitochondrial read counts, principal component analysis was applied based on highly variable genes expressed in the sequenced cells. To correct for batch effects, we integrated data from different projects using the harmony algorithm[19]. Cell populations were shown by a unified UMAP embedding space based on the harmony-corrected principal components, followed by graph-based clustering and annotation of each cluster with the respective markers (Fig. 1A). Based on the annotation results, a higher number of macrophages and lower number of CD4- and CD8-positive T cells were observed in mutant patients than in WT patients (Fig. 1B, C). Further detailed analysis showed that at the subpopulation level, exhausted CD8^+^ T cells and C1QC^+^ macrophages exhibited the highest preference for *BRAF* mutant tumors (Fig. 1D, E). Interestingly, memory CD8^+^ T cells (CD8-Tem-GZMK) decreased in *BRAF* mutant tumors (Fig. 1D, E). These results indicated that the suppressive status of CD8^+^ T cells was enriched in *BRAF* mutant CRC tumors.

**Figure 1.**
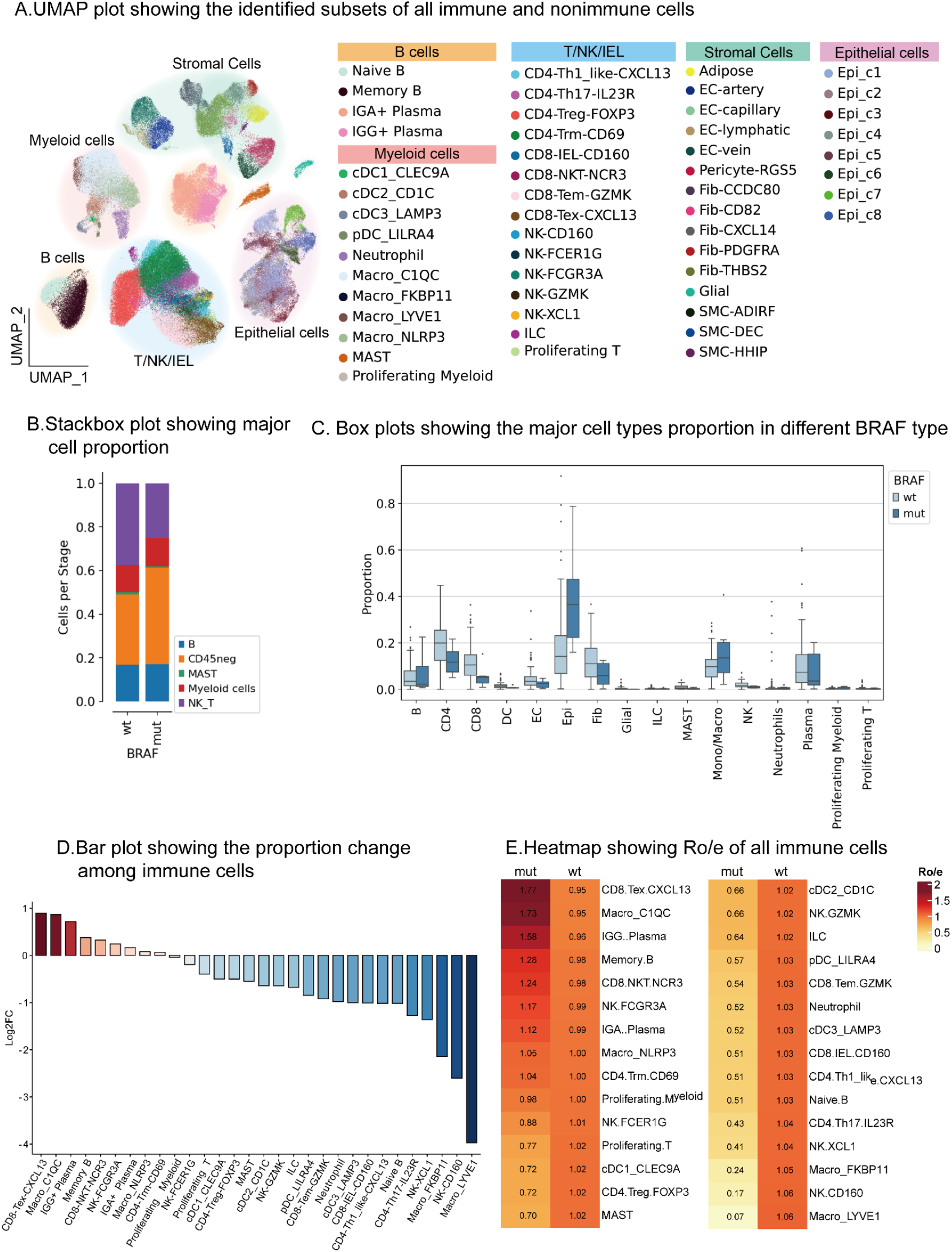
Profiling of tumor microenvironment of *BRAF* mutant colon cancer A. UMAP plot showing integrated clustering of single-cell transcriptomes from *BRAF* wild-type and mutant BRAF CRC patients. B. Comparison of cell type composition between WT and MUT tumors. C. Proportions of major immune cell types according to *BRAF* status. D. Bar plot shows the proportion of changes among immune cells. E. Heatmap showing the Ro/e of all immune sub-clusters.

### Imbalance of CD8^+^ T cell differentiation is induced in *BRAF* mutant CRC

CD8^+^ T cells act as key players in anti-cancer immunity, while terminal differentiation into exhaustion leads to CD8^+^ T cell dysfunction and cancer progression[20]. Our data showed that exhausted CD8^+^ T cells were the most abundant population in *BRAF* mutant tumors, while memory CD8^+^ T cells were decreased. These phenomena prompted us to hypothesize that the differentiation of CD8^+^ T cells may be diverted into terminal differentiation in *BRAF* mutant tumors. To test this hypothesis, we systematically compared the differential features of tumor-infiltrating CD8^+^ T cells between mutant and WT CRC. Pseudotime analysis revealed progressive exhaustion of CD8^+^ T cells in the tumor microenvironment, manifested by gradual downregulation of memory genes (GZMA, GZMK, TNF) and concurrent upregulation of immunosuppressive genes (CTLA4, CXCL13, HAVCR2, PDCD1, TIGIT, TNFRSF9) (Fig. 2A, 2B). Critically, CD8^+^ T cells in memory status were lower in mutant tumors, whereas terminally differentiated cells were higher (Fig. 2B). Indeed, the expression of memory genes was lower and exhaustion genes were higher in mutant tumors (Fig. 2C), indicating an imbalance in memory and exhaustion in mutant tumors. Specific gene expression levels of exhaustion (TOX, PDCD1, TIGIT, and TNFRSF9) were higher in mutant tumors, and memory genes were lower (Fig. 2C, 2D).

**Figure 2.**
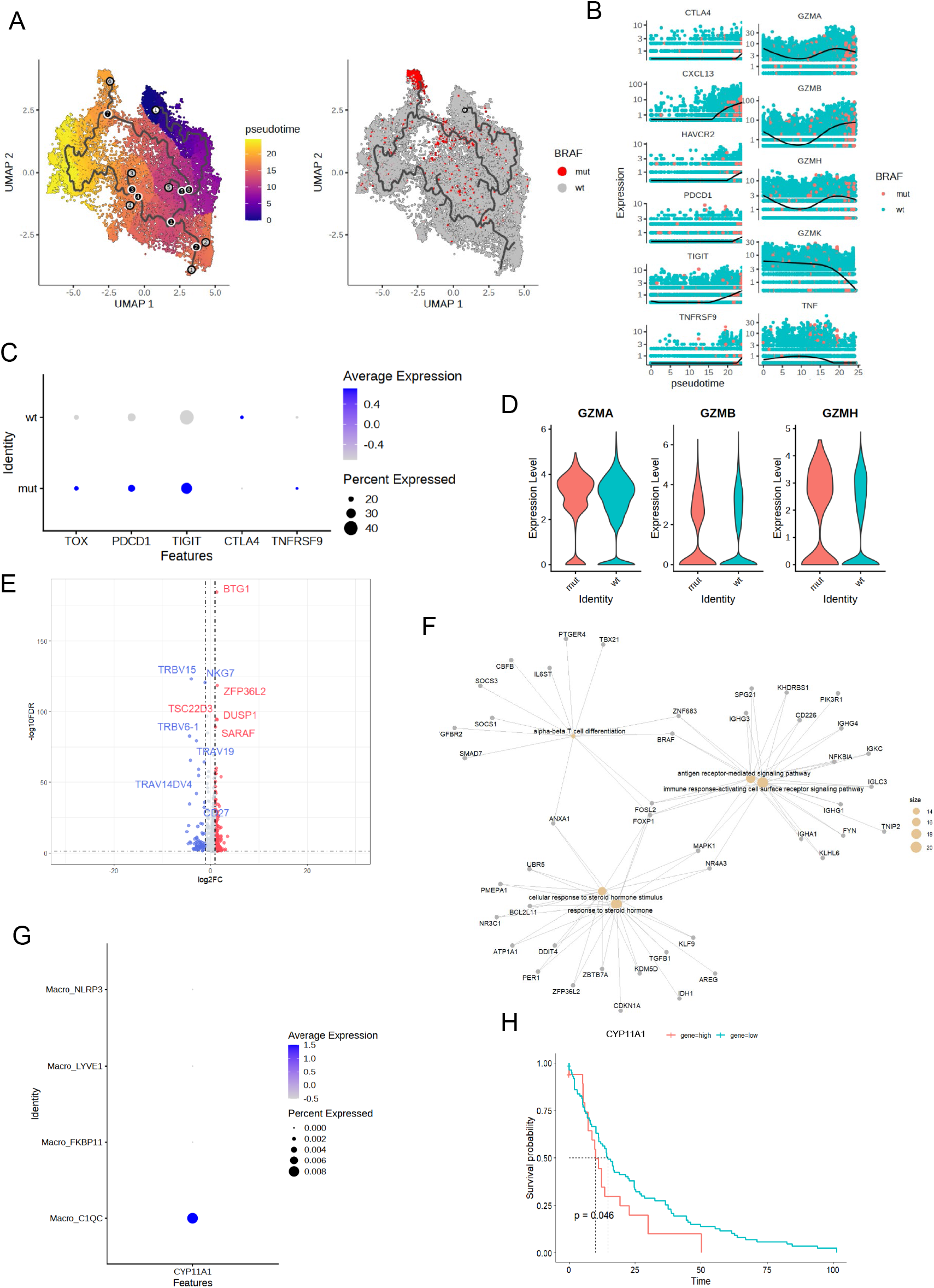
High exhaustion of CD8^+^ T cells in *BRAF* mutant colon cancer A. Pseudotime trajectory of CD8^+^ T cells showing differentiation from memory to exhausted states. B. Expression dynamics of memory (GZMK, TNF) and exhaustion (PDCD1, TIGIT) genes along the pseudotime. C. Dot plot comparing exhaustion genes expression between WT and MUT tumors. D. Violin plots showing functional gene expression between WT and MUT tumors. E. Volcano plot of differentially expressed genes in CD8^+^ T cells between *BRAF* mutant and wild-type samples. F. Gene ontology network showing functional clusters of differentially expressed genes. G. Expression of steroidogenic enzymes reveals CYP11A1 enrichment in C1QC^+^ macrophages. H. Kaplan-Meier survival analysis showing poor prognosis associated with high CYP11A1 expression in CRC.

Next, we investigated the potential mechanism of this imbalance by analyzing alterations in whole gene expression. Interestingly, in addition to the differentiation and activation pathways, the cellular response to steroid hormones was significantly increased in mutant tumors (Fig. 2E-2F). Deeper analysis revealed significantly elevated expression of CYP11A1, a rate-limiting enzyme in steroid synthesis, within the MUT tumor microenvironment, predominantly localized in C1QC^+^ macrophage subsets (Fig. 2G)[21]. Clinical prognostic analysis further confirmed that high CYP11A1 expression was significantly associated with shorter overall survival in patients with CRC (Fig. 2H). These findings suggest that *BRAF* mutant colorectal cancers harbor an immunosuppressive microenvironment characterized by increased CD8^+^ T cell exhaustion, which may reduce the immune system’s ability to kill tumors.

### High resolution spatial study identifies detailed TME of *BRAF* mutant colon cancer

Next, we investigated the spatial tumor microenvironment that participates in immunosuppressive *BRAF*-mutant CRC by spatial transcriptome profiling. To precisely delineate the composition of the immune microenvironment and intercellular relationships in *BRAF*-mutant colorectal cancers, we collected tumor specimens and performed 10X Visium HD spatial transcriptomic profiling, achieving high-resolution 2-μm spatial data (Fig. 3A). This unprecedented resolution enabled the high-fidelity capture of the tumor tissue architecture (Fig. 3B). Through histopathological image segmentation of high-definition H&E staining and Gaussian fitting-based RNA capture optimization, we reconstructed 2-μm bins into single cell-equivalent spatial spots (Fig. 3C). Subsequent dimensionality reduction and clustering of these spots revealed a comprehensive intratumoral cellular architecture (Fig. 3D-3E). Cross-modality integration was implemented by transferring cell-type annotation labels from single-cell RNA-sequencing datasets to spatial coordinates (Fig. 3F). Spatial organization analysis identified distinct tumor microenvironment domains exhibiting specific cellular interaction patterns (Fig. 3G).

**Figure 3.**
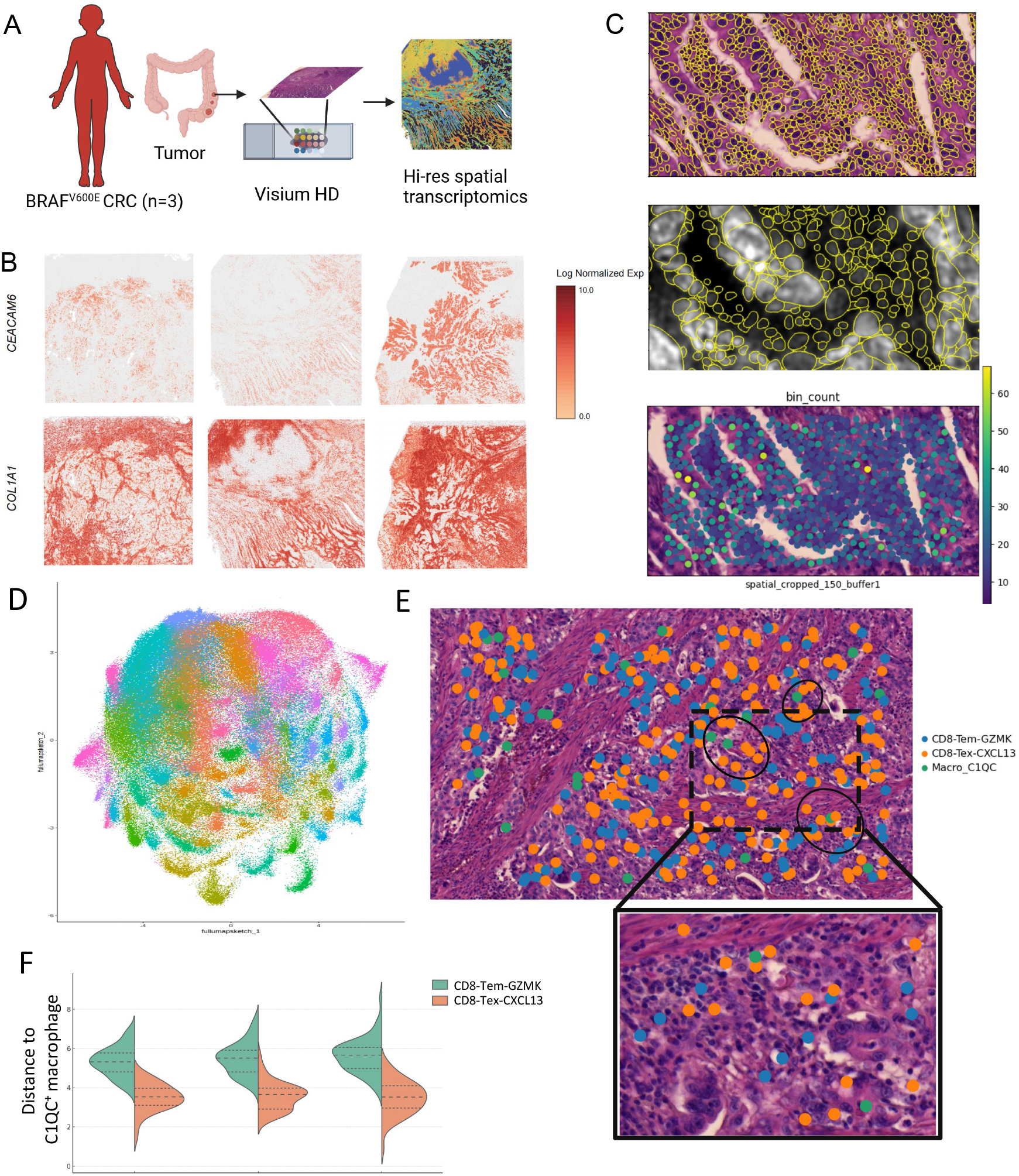
High-resolution spatial study identifies TME of *BRAF* mutant colon cancer A. Study design illustrating spatial transcriptomic workflow and sample processing. B. Tissue architecture captured by H&E staining and RNA capture features. C. Image segmentation and single-cell bin aggregation strategy for spatial deconvolution. D. UMAP showing dimensionality reduction and clustering of spatially resolved spots. E. Spatial annotation of cell showing localization of C1QC^+^ macrophage, effect memory CD8^+^ T and exhausted CD8^+^ T. F. Violin plot shows the distance to C1QC^+^ macrophage of CD8^+^ T.

### Ligand-receptor cross-talk between C1QC^+^ macrophage and CD8^+^T cells

To study the interaction of exhausted CD8^+^ T cells, we performed a systematic ligand-receptor (LR) interaction analysis, focusing on CD8^+^ T cell dynamics. Strikingly, C1QC^+^ macrophages, co-enriched with exhausted CD8^+^ T cells in mutant tumors, exhibited intensive crosstalk with CD8^+^ T populations (Fig. 4A), with significant amplification of interaction strength in MUT tumors (Fig. 4B). Spatial enrichment analysis confirmed significant colocalization between C1QC^+^ macrophages and exhausted CD8^+^ T-cells (Fig. 4C).

**Figure 4.**
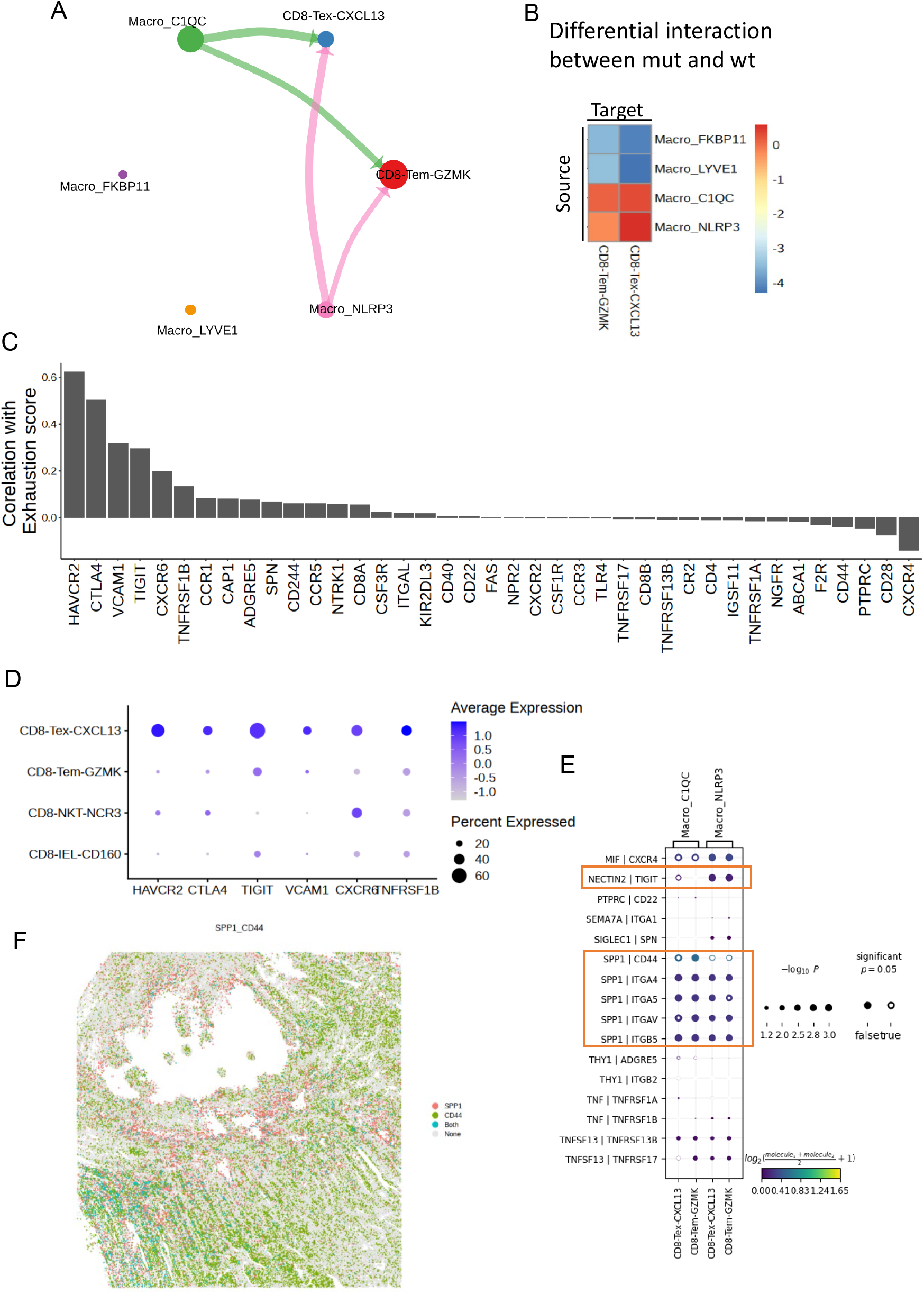
C1QC^+^ macrophage interacts with CD8^+^ T cells promoting exhaustion A. Circle plot showing ligand-receptor interaction network between C1QC^+^ macrophages and CD8^+^ T cells. B. Quantification of ligand-receptor interaction strength in *BRAF* mutant versus wild-type tumors. C. Bar plot showing the correlation between receptor expression and exhaustion score in T cells. D. Dot plot showing the expression of key immune-suppressive receptor in CD8^+^ T cells. E. Dot plot showing ligand-receptor pairs expressed in CD8^+^ T cells and C1QC^+^ macrophages. F. Spatial distribution heatmap of NECTIN and SPP1 ligands in C1QC^+^ macrophages.

The LR network revealed that multiple receptor-ligand pairs were positively correlated with CD8^+^ T cell exhaustion levels (Fig.4D-4E). C1QC^+^ macrophages in MUT tumors overexpressed immunosuppressive ligands, including NECTIN and SPP1 (Fig. 4F), with their spatial expression gradients mirroring local T-cell dysfunction patterns (Fig. 4G).

### CYP11A1 in macrophage induces CD8^+^ T cell exhaustion through pregnenolone

Both single-cell and spatial transcriptomic data indicated a significant interaction between macrophages and CD8^+^ T cells, potentially contributing to the induction of CD8^+^ T cell exhaustion. The steroidogenesis pathway centered on CYP11A1 appears to play a pivotal role in this process. CYP11A1 catalyzes the conversion of cholesterol into pregnenolone, indicating that pregnenolone may have a potential effect on exhaustion. When pregnenolone was added to in vitro cultured murine CD8^+^ T cells, there was a marked upregulation of exhaustion-associated genes (PD1, TIM3, and LAG3) (Fig. 5A).

**Figure 5.**
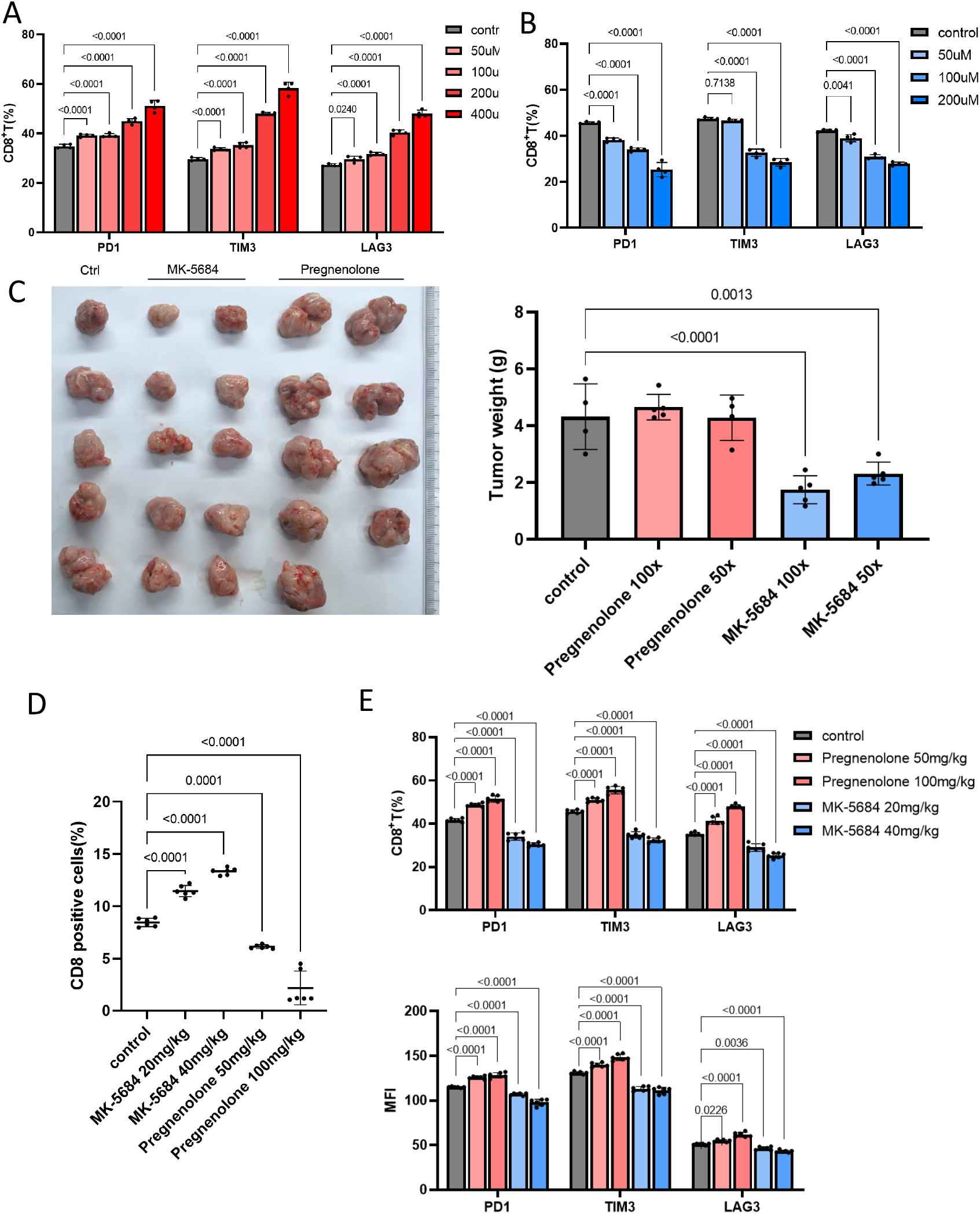
Pregnenolone induces CD8^+^ T cell exhaustion and promotes tumor progression A. Exhaustion marker expression in CD8^+^ T cells after pregnenolone treatment in vitro. B. Co-culture assay showing that CYP11A1 inhibition reduces T cell exhaustion. C. Tumor growth shows acceleration by pregnenolone and suppression by MK-5674 in vivo. D. Flow cytometry analysis showing CD8^+^ T cell proliferation is inhibited by pregnenolone and rescued by MK-5674. E. Expression of exhaustion genes (PD1, LAG3, TIM3) in tumor-infiltrating CD8^+^ T cells under different treatments.

To verify whether macrophages influence CD8^+^ T cells through pregnenolone production via CYP11A1, we established an in vitro co-culture system of macrophages and CD8^+^ T cells. Consistently, treatment with MK-5674, a selective CYP11A1 inhibitor, effectively inhibited CD8^+^ T cell exhaustion (Fig. 5B).

We further evaluated the role of the CYP11A1 pathway in tumor immunity using a murine colorectal cancer model. Exogenous administration of pregnenolone significantly accelerated tumor growth, whereas treatment with MK-5674 markedly suppressed tumor progression (Fig. 5C, 5D). A detailed analysis of tumor-infiltrating CD8^+^ T cells revealed that pregnenolone administration inhibited CD8^+^ T cell proliferation and enhanced the expression of exhaustion genes. In contrast, MK-5674 treatment promoted CD8^+^ T cell proliferation and reduced the expression of exhaustion markers (Fig. 5E–5G). Collectively, these findings demonstrate that pregnenolone production from macrophages enhances CD8^+^ T cell exhaustion, resulting in impaired anti-tumor immunity.

### C1QC^+^ macrophage-driven CD8^+^ T cell exhaustion promotes progression of *BRAF* mutant CRC and other types of cancer

Through the analysis of public transcriptomic datasets, we validated significantly elevated C1QC expression in *BRAF*-mutant colorectal cancers (Fig. 6A). Using a gene signature derived from the top 10 expressed genes in C1QC^+^ macrophages (Fig. 6B), we demonstrated that high macrophage enrichment scores were strongly correlated with reduced overall survival in patients with CRC (Fig. 6V).

**Figure 6.**
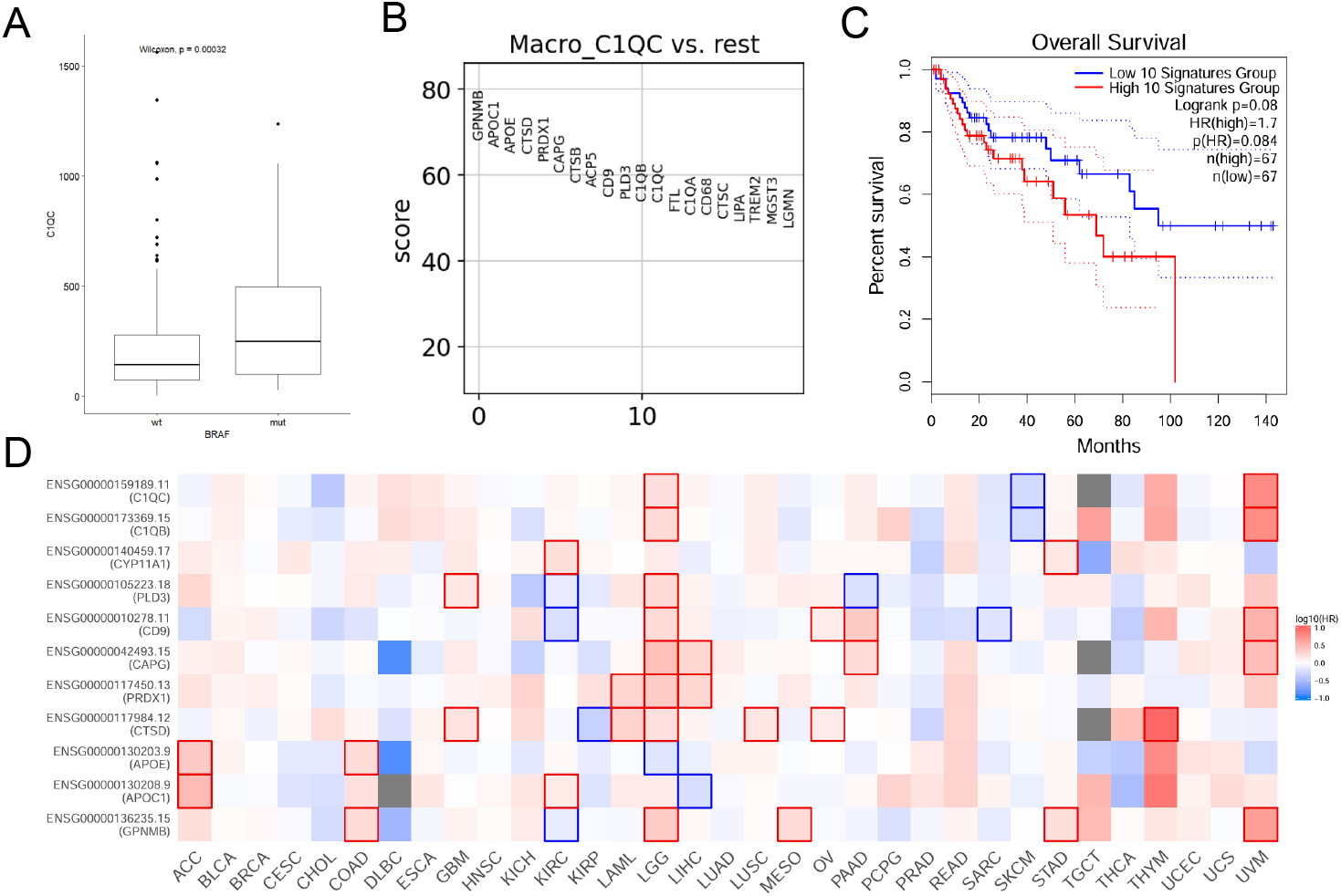
C1QC^+^ macrophage-driven exhaustion correlates with CRC prognosis and pan-cancer immunosuppression Expression of C1QC across CRC subtypes elevated in *BRAF* mutant tumors, using. Top 10 genes characterizing C1QC^+^ macrophages derived from single-cell data. C. Kaplan-Meier survival curve showing worse prognosis in CRC patients with high C1QC^+^ macrophage signature scores. D. Pan-cancer analysis showing correlation of C1QC^+^ macrophage enrichment with survival in other cancer types.

Notably, this C1QC^+^ macrophage signature showed pan-cancer prognostic significance, exhibiting similar survival correlations in brain lower-grade glioma, thymoma, and uveal melanoma (Fig. 6D). These multi-omics findings suggest a conserved mechanism whereby C1QC^+^ macrophages drive cancer progression by impairing CD8^+^ T cell functionality and promoting T cell exhaustion, thereby establishing their broad applicability as therapeutic targets across malignancies.

## Discussion

In this study, we employed high-resolution spatial analysis and single-cell RNA sequencing to map the TME of *BRAF* mutant CRC. Our findings revealed a distinct immunosuppressive landscape characterized by the enrichment of exhausted CD8^+^ T cells and C1QC^+^ macrophages. This high-resolution spatial and cellular characterization provides a comprehensive view of the TME in *BRAF* mutant CRC, highlighting the critical role of cellular interactions in shaping the immune response.

A key link between exhausted CD8^+^ T cells and C1QC^+^ macrophages is CYP11A1 and the response to steroid hormones in *BRAF* mutant tumors. CYP11A1, a mitochondrial enzyme responsible for converting cholesterol to pregnenolone, has been implicated in promoting an immunosuppressive state through steroidogenesis[22]. Our data show that CYP11A1 high expression in C1QC^+^ macrophages correlates with increased CD8^+^ T cell exhaustion, suggesting a direct link between pregnenolone synthesis and immune evasion. Pregnenolone and its derivatives have been primarily studied in the context of hormone-dependent malignancies, where they act as upstream regulators of sex steroid signaling. Traditionally, steroidogenesis has been found to be relevant in prostate or breast cancer[23]. In our study, both in vitro and in vivo investigations demonstrated that pregnenolone drives immune evasion by promoting CD8^+^ T cell exhaustion. Therefore, our work reveals that CYP11A1-expressing C1QC^+^ macrophages induce CD8^+^ T cell exhaustion via pregnenolone synthesis in *BRAF* mutant CRCs. Our study expands the oncologic relevance of this metabolic pathway by demonstrating that pregnenolone synthesis also plays a critical role in non-endocrine tumors such as colorectal cancer.

Our results suggest that intratumoral steroid production, particularly pregnenolone production, may represent a conserved immunosuppressive strategy used across diverse malignancies. This is further supported by the pan-cancer prognostic relevance of C1QC^+^ macrophage signatures in gliomas, melanomas, and thymomas.

Pregnenolone metabolism is a novel therapeutic target. Indeed, our study found that targeting CYP11A1 effectively restored the anti-tumor function of CD8^+^ T cells.

These insights warrant further investigation of steroid biosynthesis pathways in other cancer types, and suggest that reprogramming the metabolic-immune axis of the TME could offer a promising new avenue for cancer treatment. Together, these findings open new avenues for therapeutic intervention and highlight the potential of targeting the pregnenolone pathway to enhance anti-tumor immunity.

## Materials and Methods

### Ethical approval

All experimental protools involving human subjects and biological samples were performed in accordance with ethical standards established by the Committee of Tsinghua University. The acquisition and use of clinical specimens were approved by the Ethics Committee at Peking Union Medical College Hospital. Written consent was obtained from all donors prior to participation. The study design complied with the regulations of the Association for Assessment and Accreditation of Laboratory Animal Care International and was formally sanctioned by the Institutional Animal Care and Use Committee (IACUC) at Tsinghua University following rigorous review. Specifically, in murine tumor models, dimensional parameters of neoplasms were strictly maintained within the ethical boundaries prescribed by the IACUC, ensuring that tumor volumes remained below the 2 cm^3^ threshold throughout the investigation period.

### Clinical sample collection and preparation

All the enrolled patients were histopathologically diagnosed with *BRAF* mutant colorectal adenocarcinoma. Patients with autoimmune diseases, prior malignancies, or a history of preoperative chemotherapy, radiotherapy, or immunotherapy were also excluded. Tumor tissues were collected within 15 min of surgical excision, following thorough clearance of the blood and necrotic tissue. Comprehensive pathological profiles of the patients are detailed in Supplementary Table 1. All specimen collections were performed at Peking Union Medical College Hospital.

### Animals

Six-to eight-week-old C57BL/6 mice were purchased from Vital River Laboratory Animal Technology and used for CD8+ T cell isolation. Six-to eight-week-old OT-I mice were purchased from The Jackson Laboratory and used for the isolation of OT-I CD8+ T cells. All mice were maintained in pathogen-free facilities and used strictly in accordance with protocols approved by the IACUC of Tsinghua University. The study complied with all relevant ethical regulations regarding animal research. Sex was not considered in the study design and analysis because this study was not designed to detect sex differences. Only male mice were used in this study, and all mouse data were collected from male mice.

### Cell lines

The MC38-OVA cell line was obtained from P. Jiang Laboratory (Tsinghua University, Beijing, China). 293T cells were purchased from the American Type Culture Collection (ATCC; CRL-3216). MC38-OVA and 293T cells were cultured in DMEM medium (DMEM; CM10013). Primary CD4+ and CD8+ T cells were cultured in RMPI-1640 medium (macgene, CM10041). All the cell lines were authenticated and tested for mycoplasma contamination. None of the commonly misidentified cell lines were used in this study. Furthermore, we authenticated all cell lines using a short tandem repeat profiling method according to a reported protocol.

### Isolation and stimulation of primary CD8^+^ T cells

Mouse CD8^+^ T cells were isolated from mouse spleens using a biotin selection kit. Unless otherwise indicated, mouse CD8^+^ T cells were stimulated with 3.5 µg/mL anti-CD3 and 1 µg/mL anti-CD28 antibodies. The OT-I CD8^+^ T cells from OT-I mice were stimulated with 1 µg/mL OVA_257–264_ (SIINFEKL) peptide. All CD8^+^ T cells were cultured in RMPI-1640 medium (macgene, CM10041) with 100 U/mL IL-2. For *in vitro* exhaustion, CD8^+^ T cells were stimulated 4 times with 2 days’ interval.

### In Vitro Pregnenolone Treatment of CD8^+^ T Cells

CD8+ T cells were plated in 24-well plates (1 × 10^6 cells/well) in RPMI-1640 medium supplemented with 10% FBS. For stimulation, pregnenolone (Heowns, P-66146) was added to a final concentration of 100 μM and incubated for 48 hours. Expression of exhaustion markers (PD1, TIM3, LAG3) was assessed by flow cytometry.

### Co-culture of CD8^+^ T Cells with BMDMs and CYP11A1 Inhibition

Bone marrow-derived macrophages (BMDMs) were generated by culturing bone marrow cells in the presence of 20 ng/mL M-CSF and for 7 days. MK-5684 (Opevesostat; TargetMol, T77627), a selective CYP11A1 inhibitor, was added at a final concentration of 100 nM. Mature BMDMs were co-cultured with CD8+ T cells at a 1:2 ratio (macrophage:T cell) to investigate its effect on T cell exhaustion. Co-cultures were maintained for 72 hours before phenotypic and gene expression analyses.

### In Vivo Pregnenolone Treatment

C57BL/6 mice were subcutaneously inoculated with 5 × 10^5 MC38-OVA tumor cells. Pregnenolone was dissolved in 0.5% SDS and then resuspended in peanut oil to a final dose of 60 mg/kg. Mice received intraperitoneal injections every other day, starting from day 4 post-tumor inoculation. Tumor volumes were measured every two days using digital calipers.

### In Vivo MK-5684 Treatment

MK-5684 was suspended in 0.5% Tween and 0.5% aqueous methyl cellulose. Mice received oral gavage of MK-5684 at 20 mg/kg twice daily for seven consecutive days. Tumor growth was monitored throughout the treatment period. At endpoint, tumors were harvested and tumor-infiltrating lymphocytes were analyzed via flow cytometry to evaluate the immune status of CD8^+^ T cells.

### Dissociation of tumor-infiltrating cells from primary tissues

Mouse tumor samples were enzymatically digested into single cells. Then, samples were added with collagenase IV and shaked at 37^°^C for 1.5 hours for digestion. The digested samples were filtered through a single-cell strainer to remove tissue debris. After washing, the samples were ready for subsequent flow cytometry analysis. For cytokine analysis, samples were treated with anti-CD3, anti-CD28 antibodies and BFA/Monensin for 5 hours prior flow cytometry analysis.

### Flow cytometry

For surface staining, cells were harvested and stained with antibodies at the recommended dilution prior to Flow Cytometry analysis. For intracellular proteins including cytokines, cells were harvested and washed, fixed with eBioscience™ IC Fixation Buffer (00-8222-49), permeabilized with eBioscience™ Permeabilization Buffer (00-8333-56) and stained according to the manufacturer protocol. For intranuclear proteins, cells were harvested and fixed with eBioscience™ Fixation/Permeabilization Diluent and Concentrate (00-5223-56 and 00-5123-43), permeabilized with eBioscience™ Permeabilization Buffer (00-8333-56) and stained. Stained cells were washed prior to FACS analysis. BD FACSuite Flow Cytometry Software was used for FACS signal collection. Positive events were determined by isotype control gating for each antibody. Cells were first gated using FSC/SSC characteristics to exclude debris, followed by gating FSC-W and FSC-H, then SSC-W and SSC-H to eliminate nonsinglets. Then target cells were gated the population of interest by specific stain. Data analysis was carried out using FlowJo.

### Single-Cell RNA Sequencing Data Acquisition and Processing

Publicly available scRNA-seq datasets of colorectal cancer were retrieved from Synapse(syn26844071) and annotated using whole-exome sequencing data to classify samples into BRAF mutant and wild-type groups. After standard quality control (excluding cells with >25% mitochondrial reads and <200 or >7000 genes), a total of 191,292 cells were retained. Batch corrections were performed using a harmony algorithm. Clustering and UMAP visualizations were conducted using Seurat v5.2. The cell types were annotated based on canonical markers.

### Trajectory and Gene Expression Analysis of CD8^+^ T Cells

CD8 + T cell subsets were re-clustered and analyzed using Monocle 2 to construct pseudo-time trajectories. Differential expression analysis was performed to identify genes associated with exhaustion and memory. Gene set enrichment analysis (GSEA) was conducted using clusterProfiler and curated gene sets from KEGG and other literature sources.

### Spatial Transcriptomics

High-resolution spatial transcriptomic profiling was performed on BRAF mutant tumor samples using 10x Genomics Visium HD technology. The tissue sections were processed according to the manufacturer’s protocols. RNA capture spots were reconstructed at single-cell resolution using Bin2Cell. Cell-type annotations from the scRNA-seq data were transferred using label transfer algorithms. Cell-cell spatial colocalization and interaction domains were defined via correlation analysis and clustering.

### Cell-Cell Communication and Ligand-Receptor Analysis

CellChat v2.1.2 was used to infer ligand-receptor interactions between C1QC^+^ macrophages and CD8 + T cells from scRNA-seq data. Interactions were filtered based on expression confidence and cell number thresholds. Spatial co-expression and interaction strength were validated using Visium data through permutation-based colocalization scoring.

### Survival analysis

Kaplan-Meier survival curves were generated and statistically analyzed using ggsignif 0.6.4 (). Survival differences between groups were assessed using log-rank tests, with p-values < 0.05 considered statistically significant.

### Other statistical analyses

For bioinformatic analysis, group comparisons were analyzed using a two-sided unpaired Student’s t-test, with statistical significance defined as p < 0.05. For experimental data analysis, quantitative measurements, displayed as mean ± SD, were obtained from at least triplicate biological replicates. Statistical contrasts between groups were analyzed using a two-sided unpaired Student’s t-test, with statistical significance defined as p < 0.05. The normality assumption for the t-test was adopted without formal verification of data distribution. All raw data points are displayed as graphical representations. Experiments were designed with three–ten replicates per independent group or condition. Biological duplicates and statistical analyses have been provided in the figure legends. Sample sizes were not predetermined using statistical methods but were aligned with commonly adopted parameters in comparable studies. No experimental data were omitted from statistical evaluations. In the absence of specific statements, the experiments were conducted without randomization, and the researchers were aware of the allocation during both the experiments and outcome assessment. Investigators remained masked to the group assignments throughout the animal experimentation data recording, non-animal experimental procedures, and analytical evaluations. The minimum cohort size (n ≥5) of the animals was based on feasibility and specimen accessibility to validate the findings.

## Resource availability

### Lead contact

For additional details or resource inquiries, please contact the corresponding author

### Materials availability

No novel reagents were synthesized in this study.

### Data and code availability

This study employs established analytical methods without the development of novel algorithms. The R and Python codes supporting the key analyses are available upon request from the corresponding author.

## Acknowledgments

We thank Prof. Cheng Li for bioinformatics help and Yan Liu and the Tsinghua University Branch of China National Center for Protein Sciences (Beijing) and Tsinghua University Technology Center for Flow cytometry support. This study was supported by the CAMS Innovation Fund for Medical Sciences (CIFMS 2023-I2M-3-012).

## Author contributions

Y.B. and J.C. designed the study. Zihao Zhao performed all analyses and experiments under the supervision of J.C. X.W., and L.L. performed patient sample collection and clinical analysis under the supervision of Y.B. Y.B. and J.C. supervised the research. Zihao Zhao wrote the manuscript. All authors have commented on the manuscript.

## Reference

[1] Sung H, Ferlay J, Siegel RL, Laversanne M, Soerjomataram I, Jemal A, Bray F: Global Cancer Statistics 2020: GLOBOCAN Estimates of Incidence and Mortality Worldwide for 36 Cancers in 185 Countries. CA: a cancer journal for clinicians 2021, 71(3):209–249.

[2] Guo L, Wang Y, Yang W, Wang C, Guo T, Yang J, Shao Z, Cai G, Cai S, Zhang L et al: Molecular Profiling Provides Clinical Insights Into Targeted and Immunotherapies as Well as Colorectal Cancer Prognosis. Gastroenterology 2023, 165(2):414-428.e417.

[3] Xu T, Wang X, Wang Z, Deng T, Qi C, Liu D, Li Y, Ji C, Li J, Shen L: Molecular mechanisms underlying the resistance of BRAF V600E-mutant metastatic colorectal cancer to EGFR/BRAF inhibitors. Therapeutic advances in medical oncology 2022, 14:17588359221105022.

[4] García-Alfonso P, Lièvre A, Loupakis F, Tadmouri A, Khan S, Barcena L, Stintzing S: Systematic review of randomised clinical trials and observational studies for patients with RAS wild-type or BRAF(V600E)-mutant metastatic and/or unresectable colorectal cancer. Critical reviews in oncology/hematology 2022, 173:103646.

[5] Corcoran RB, Ebi H, Turke AB, Coffee EM, Nishino M, Cogdill AP, Brown RD, Della Pelle P, Dias-Santagata D, Hung KE et al: EGFR-mediated re-activation of MAPK signaling contributes to insensitivity of BRAF mutant colorectal cancers to RAF inhibition with vemurafenib. Cancer discovery 2012, 2(3):227–235.

[6] Corcoran RB, André T, Atreya CE, Schellens JHM, Yoshino T, Bendell JC, Hollebecque A, McRee AJ, Siena S, Middleton G et al: Combined BRAF, EGFR, and MEK Inhibition in Patients with BRAF(V600E)-Mutant Colorectal Cancer. Cancer discovery 2018, 8(4):428–443.

[7] Tabernero J, Grothey A, Van Cutsem E, Yaeger R, Wasan H, Yoshino T, Desai J, Ciardiello F, Loupakis F, Hong YS et al: Encorafenib Plus Cetuximab as a New Standard of Care for Previously Treated BRAF V600E-Mutant Metastatic Colorectal Cancer: Updated Survival Results and Subgroup Analyses from the BEACON Study. Journal of clinical oncology : official journal of the American Society of Clinical Oncology 2021, 39(4):273–284.

[8] Wang D, Wang L, Zhang W, Xu K, Chen L, Guo Z, Wu K, Huang D, Zhao Y, Yao M et al: Extracellular vesicle-mediated gene therapy targets BRAF(V600E)-mutant colorectal cancer by inhibiting the MEK1/2-ERK1/2 pathway. Journal of nanobiotechnology 2025, 23(1):129.

[9] Sun C, España S, Buges C, Layos L, Hierro C, Manzano JL: Treatment of Advanced BRAF-Mutated Colorectal Cancer: Where We Are and Where We Are Going. Clinical colorectal cancer 2022, 21(2):71–79.

[10] Poulikakos PI, Sullivan RJ, Yaeger R: Molecular Pathways and Mechanisms of BRAF in Cancer Therapy. Clinical cancer research : an official journal of the American Association for Cancer Research 2022, 28(21):4618–4628.

[11] Giménez N, Martínez-Trillos A, Montraveta A, Lopez-Guerra M, Rosich L, Nadeu F, Valero JG, Aymerich M, Magnano L, Rozman M et al: Mutations in the RAS-BRAF-MAPK-ERK pathway define a specific subgroup of patients with adverse clinical features and provide new therapeutic options in chronic lymphocytic leukemia. Haematologica 2019, 104(3):576–586.

[12] Hornsteiner F, Vierthaler J, Strandt H, Resag A, Fu Z, Ausserhofer M, Tripp CH, Dieckmann S, Kanduth M, Farrand K et al: Tumor-targeted therapy with BRAF-inhibitor recruits activated dendritic cells to promote tumor immunity in melanoma. Journal for immunotherapy of cancer 2024, 12(4).

[13] Erkes DA, Cai W, Sanchez IM, Purwin TJ, Rogers C, Field CO, Berger AC, Hartsough EJ, Rodeck U, Alnemri ES et al: Mutant BRAF and MEK Inhibitors Regulate the Tumor Immune Microenvironment via Pyroptosis. Cancer discovery 2020, 10(2):254–269.

[14] Roy S, Sipthorp J, Mahata B, Pramanik J, Hennrich ML, Gavin AC, Ley SV, Teichmann SA: CLICK-enabled analogues reveal pregnenolone interactomes in cancer and immune cells. iScience 2021, 24(5):102485.

[15] Mahata B, Pramanik J, van der Weyden L, Polanski K, Kar G, Riedel A, Chen X, Fonseca NA, Kundu K, Campos LS et al: Tumors induce de novo steroid biosynthesis in T cells to evade immunity. Nature communications 2020, 11(1):3588.

[16] Ge S, Cheng D, Zhang X, Xu T, Wang Z, Dong F, Su L, Song J, Wang J, Li J et al: Using genotype to assist clinical surveillance: a retrospective study of Chinese familial adenomatous polyposis patients. American journal of cancer research 2022, 12(9):4254–4266.

[17] El Hajj Y, Shahin T, Dieng MM, Alshaikh M, Khair M, Manikandan V, Idaghdour Y: Pregnenolone sulfate induces transcriptional and immunoregulatory effects on T cells. Scientific reports 2024, 14(1):6782.

[18] Joanito, I., Wirapati, P., Zhao, N. et al. Single-cell and bulk transcriptome sequencing identifies two epithelial tumor cell states and refines the consensus molecular classification of colorectal cancer. Nat Genet 54, 963–975 (2022).

[19] Korsunsky I, Millard N, Fan J, et al. Fast, sensitive and accurate integration of single-cell data with Harmony. Nature Methods. 2019.

[20] Wherry EJ, Kurachi M. Molecular and cellular insights into T cell exhaustion. Nature Reviews Immunology. 2015.

[21] Li X, Xu J, Yang Y, et al. CYP11A1 is a novel diagnostic and prognostic biomarker in multiple human cancers. BMC Cancer. 2021.

[22] Endogenous Glucocorticoid Signaling Regulates CD8+ T Cell Differentiation and Development of Dysfunction in the Tumor Microenvironment, Acharya, Nandini et al. Immunity, Volume 53, Issue 3, 658-671.e6

[23] Lee, J.JK., Jung, Y.L., Cheong, TC. et al. ERα-associated translocations underlie oncogene amplifications in breast cancer. Nature 618, 1024–1032 (2023). 10.1038/s41586-023-06057-w

